# Mathematical modeling supports fate restriction in neurogenic progenitors of the embryonic ventral spinal cord

**DOI:** 10.1101/384347

**Authors:** Manon Azaïs, Eric Agius, Stéphane Blanco, Jacques Gautrais, Angie Molina, Fabienne Pituello, Jean-Marc Tregan

## Abstract

In the developing neural tube in chicken and mammals, neural stem cells proliferate and differentiate according to a stereotyped spatio-temporal pattern. Several actors have been identified in the control of this process, from tissue-scale morphogens patterning (Shh, BMP) to intrinsic determinants in neural progenitor cells. In a previous study (Bonnet et al. eLife 7, 2018), we have shown that the CDC25B phosphatase promotes the transition from proliferation to differentiation in a cell-cycle independent fashion. In this study, we set up a mathematical model linking progenitor modes of division to the dynamics of progenitors and differentiated populations. Here, we build on this previous model to propose a complete dynamical picture of this process. We start from the standard model in which progenitors are homogeneous and can perform any type of divisions (proliferative division yielding two progenitors, asymmetric neurogenic divisions yielding one progenitor and one neuron, and terminal symmetric divisions yielding two neurons). We constraint this model using published data about mode of divisions and population dynamics of progenitors/neurons at different developmental stages (Saade et al. Cell Reports 4, 2013), and check the effect of CDC25B gain of function in this context. Next, we explore the scenarios in which progenitors population is actually split into two different pools, one of which composed of cells that have lost the capacity to perform proliferative divisions (fate restriction). We show that one such scenario appears relevant and calls for further identification of the alternative role of CDC25B in such a fate restriction.

## 1 Introduction

How can a small number of apparently initially homogeneous neural stem cells (NSCs) give rise to the tremendous diversity of differentiated neurons and glia found in the adult central nervous system (CNS) ? The long-standing paradigm just claims: by proliferating first, and then restricting the kind of cells a progenitor can produce given its situation in time and space. How the progenitors fate progression occurs in different contexts is still under scrutiny.

In Drosophila, NSCs are multi-potent and divide asymmetrically to generate different types of progenies in a stereotypical manner. The study of mechanisms by which a single NSC can generate a wide repertoire of neural fates in this system is in fast progress [1]. In particular, several studies have highlighted the deterministic role of a series of sequentially expressed transcription factors in the temporal specification of Drosophila NSCs [2], albeit further studies substantiated that they are possibly under the control of some extrinsic (especially nutritional) factors [3].

In the mammalian cerebral cortex, the diversity of neural progenies has been linked to different types of proliferating progenitors that have been characterized [4]. In the neural tube more specifically, two morphogen gradients (Shh and BMP) have been identified that induce NSCs to adopt different identities based on their position along the dorso-ventral axis [5, 6, 7]. This spatial patterning system ensures that different types of daughter cells are generated in an adequate stereotypical spatial order despite the fact that these progenitors are considered having initially the same potential. The molecular players that control this spatial specification and their mode of action have been characterized [6]. However, little is still known yet about how *temporal* differentiation of NSCs is orchestrated, namely what controls the timing of their transition from proliferation to differentiation at a given location [8].

Following our previous study on the role of a cell cycle regulator, the phosphatase CDC25B, in the control of neurogenesis in the chicken neural tube [9], our question here is to examine whether the progenitors that perform asymmetric neurogenic divisions have lost their proliferative power at some time point (fate restriction). From that point of view, we note that ventral neural progenitors in the neural tube have been already shown to display a fate switch, transiting from early motoneurons production to late oligodendroglial production, under the control of Shh induction [10]. Here, we consider the possibility that similar kind of switch operates sooner in the same population and sustains the transition from pure proliferative divisions to neurogenic divisions.

To give sense to this hypothesis, we start from the model of fate transition we have proposed in our previous paper about the instrumental role CDC25B plays in the progression from proliferative to neurogenic divisions [9]. In the spirit of Lander et al.[11], modeling is used here as a way to gain clarity in the face of intricacy. To this end, we build on the model we have presented in the work reported in [9]. This model considered Mode of Division (hereafter MoD) as stationary over the 24 hours of our experiment. We now consider their change over time in order to extend this model over the full dynamics of ventral spinal cord motoneurons production. This extension over time uses the data published by Marty’s team [12] who measured the two essential components of this system at different times of spinal cord development: MoD on the one hand, and dynamics of Progenitors / Neurons (P/N) populations on the other hand.

From the modeling point of view, we point out the importance of being explicit about what are the observables entities (that are experimentally measured) in this system, and what are the conceptual entities we are thinking with. Explicitly, we first propose below a “basic” model which is based on the observable entities only (MoD and P/N evolutions). We use this model to make the link between these observables entities and check how experimentally measured evolution of modes of divisions can explain the evolution of cellular populations of progenitors (P-cells) and differentiated cells (N-cells).

Next, we explore the idea of fate restriction in the progenitors. For this, we have to define two conceptual entities that implement this fate restriction hypothesis, namely we define two non observable (so far) populations of progenitors, each kind of progenitors being able to perform only a restricted set of MoD. We identified three scenarios compatible with such fate restriction. In order to check the structural consequences of each scenario, we reconstruct for each of them what should be the non observable time transitions in their MoD if we take as a constraint that they must match the observable ones. We then check for the relevance of a given scenario by comparing its predicted P/N evolutions to the observable ones.

In the end, we advocate that one scenario is of great relevance: the scenario in which a first (asymmetric) neurogenic division has a ratchet effect so that once a progenitor has produced a neuron, it looses its proliferative power.

## 2 Results

### 2.1 Basic Model for the Dynamics of Mode of Division

For the sake of clarity, we recall here the basic model we designed in Bonnet et al. [9]. We consider a population of cells at time *t*, some of which are proliferating progenitors *P*(*t*), and the others are differentiated neurons *N*(*t*). The dividing progenitors can undergo three kinds of fate, yielding:

- proliferative divisions ending with two progenitors (pp-divisions)
- asymmetric divisions ending with one progenitor and one neuron (pn-divisions)
- terminal divisions ending with two neurons (nn-divisions)

Let us denote :

*η* the rate at which P-cells undergo divisions (in fraction of the P-pool per unit time)

*α*_pp_(*t*) the fraction of dividing cells undergoing pp-divisions

*α*_pn_(*t*) the fraction of dividing cells undergoing pn-divisions

*α*_nn_(*t*) the fraction of dividing cells undergoing nn-divisions

The fractions of pp-, pn- and nn-divisions can evolve with time, under the constraint that *α_pp_*(*t*) + *α_pn_*(*t*) + *α_nn_*(*t*) = 1.

The time change *Ṗ*(*t*) of pool *P*(*t*) (resp. *Ṅ*(*t*)) is then given by the balance equation at time *t*, reading:

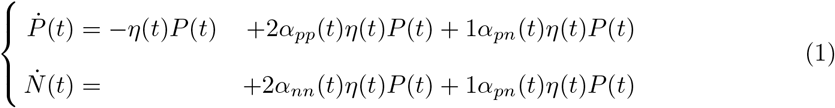

where in the first equation :

- −*η*(*t*)*P*(*t*) quantifies the rate at which P-cells disappear from the pool *P*(*t*) because they divide. The quantity of disappearing P-cells between *t* and *t* + *dt* is then *η*(*t*)*P*(*t*)*dt*
- *α_pp_*(*t*)*η*(*t*)*P*(*t*) quantifies the fraction of this quantity that undergoes a pp-division ; it doubles to yield 2 P and adds up to the pool P(t) (hence the factor 2)
- *α_pn_*(*t*)*η*(*t*)*P*(*t*) quantifies the fraction of this quantity that undergoes a pn-division ; it doubles to yield 1 P and 1 N, so only half (the P part) adds up to the pool P(t) (hence the factor 1)

Correspondingly in the second equation :

- *α_nn_*(*t*)*η*(*t*)*P*(*t*) quantifies the fraction of this quantity that undergoes a nn-division ; it doubles to yield 2 N and adds up to the pool N(t) (hence the factor 2)
- *α_pn_*(*t*)*η*(*t*)*P*(*t*) is the fraction of this quantity that undergoes a pn-division ; it doubles to yield 1 P and 1 N and only half (the N part) adds up to the pool N(t) (hence the factor 1)

System (1) is a textbook continuous-time representation of population dynamics. It is a very good approximation of the evolution of progenitors and neurons [9], considering that division events are instantaneous (M-phase is very short compared to the cell cycle duration), and occur uniformly in time (asynchronously).

The evolution of the MoD drives the evolution of the balance between proliferation and differentiation. We denote this balance:

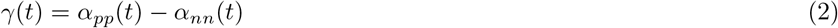

so that γ = 1 for pure proliferative process, γ = −1 for pure differentiating process, and γ = 0 for pure self-renewing process.

Using γ(*t*), the system (1) can be rewritten:

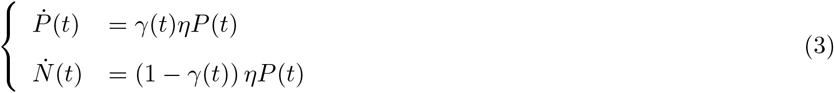

so that the general form of the solution for the evolution of the pools is given by:

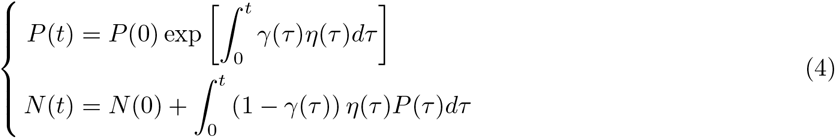

Starting from an initial configuration, *P*(0) = 1, *N*(0) =0 at time *t*_0_ and considering a steady rate *η*(*t*) = *η*, the system evolution will be only driven by the two unknown functions *α_pp_*(*t*) and *α_nn_*(*t*) that specifies *γ*(*t*).

In the embryonic spinal cord, data were collected by Saade et al.[12], that we will use to constrain the unknowns (Fig. 1a).

**Figure 1.**
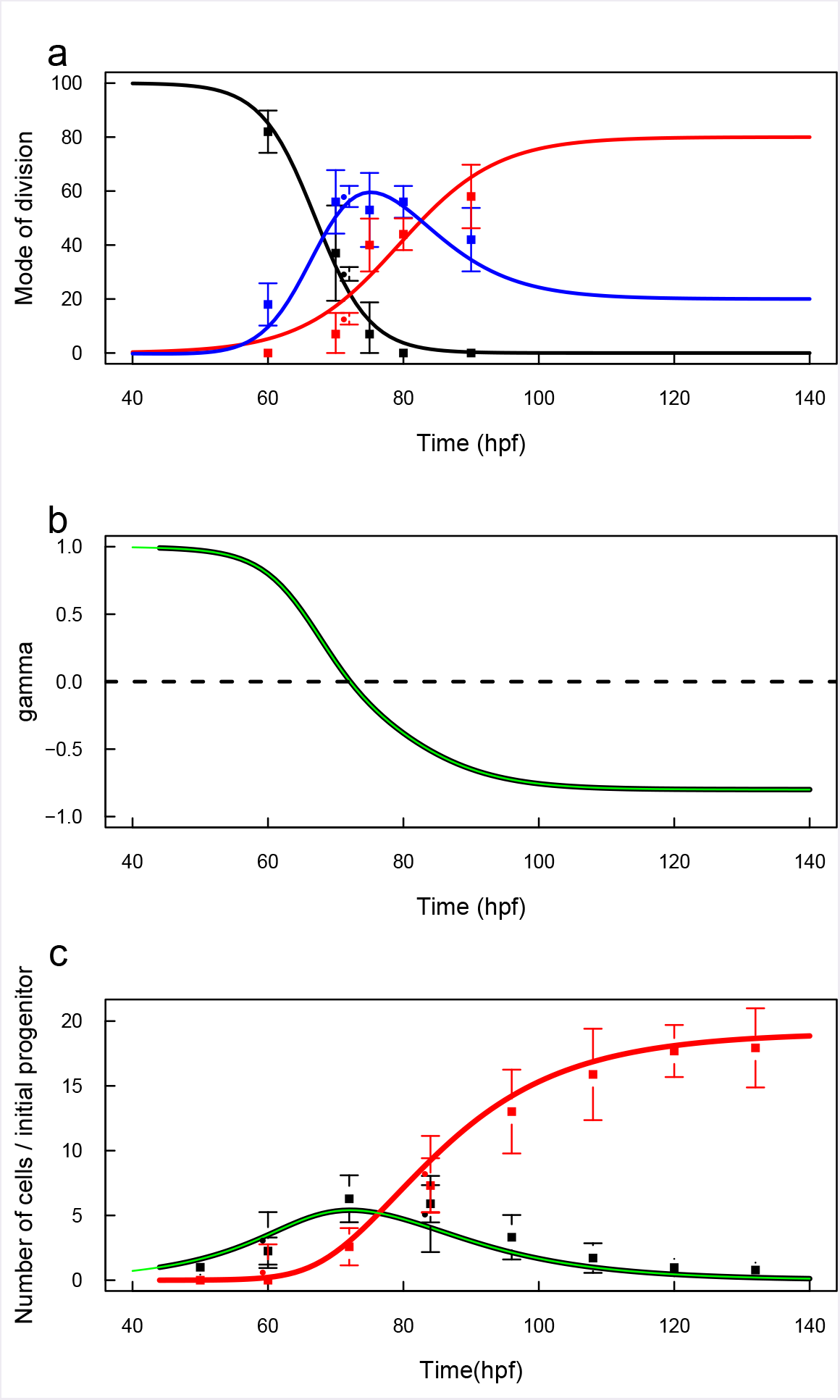
Basic Model for the dynamics of Modes of Division (MoD) in the developing ventral spinal cord. a) Discrete times estimation of MoD was retrieved from Saade et al. 2013 (square dots) and Bonnet et al. 2018 (circles), with their 95% confidence interval (bars). Black points indicate the evolution of PP-divisions, red points indicate the evolution of NN-divisions, and blue points indicate complementary (to one) PN-divisions. Smooth evolution curves were fitted separately for PP-divisions (black line) and NN-divisions (red line). Blue line fits the complementary PN-divisions, as expected. b) Corresponding evolution of the balance proliferation / differentiation γ (gamma), defined as γ(*t*) = *α_PP_*(*t*) — *α_NN_*(*t*). c) Discrete times estimation of the evolutions of the pools of progenitors (black squares) and neurons (red squares) was retrieved from Saade et al. 2013, and rescaled to correspond to the number of cells per progenitor originally present (12 progenitors at time 48 hpf). Circle points indicate estimates of P/N proportion retrieved from Bonnet et al. 2018, and scaled to the total amount of cells. Black and red lines represent respectively the predicted evolution of the P pool and the N pool, given the evolution of MoD shown in a), initial condition and population cycle time as indicated in the text. They were obtained by numerical integration of equation 4. Green line reports the analytical solution 8.

In this system, pp-divisions are largely dominant at the beginning of the process so that proliferation increases the pool of progenitors for a while, but their proportion decreases with time so that the process ends with terminal differentiation. From a minimalistic approach, we constrain the shape of the unknown functions with a minimal set of parameters for pp-divisions and nn-divisions time profiles (the pn-divisions profile being constrained to be the complement to 1). The pp-divisions time transition from *α_pp_*(*t*_0_) = 1 down to *α_pp_*(*t* → ∞) = 0 will be characterized by a characteristic time *τ_pp_* for the time of transition, with *α_pp_*(*τ_pp_*) = 0.5, and a characteristic scale *σ_pp_* indicating the sharpness of transition. A standard form for this is:

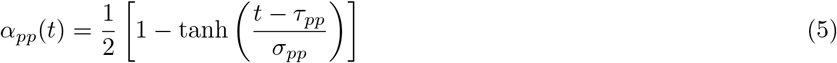

Least-square error estimation of the two parameters yields: *τ_pp_* = 67.0 hpf and *σ_pp_* = 8.0 hpf. The adjusted profile fits pretty well the data (sq. error = 0.007).

We next fitted the same kind of profile for the nn-divisions. We constrained the shape of nn-divisions evolution following:

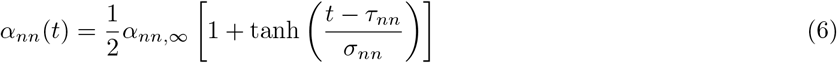

where *α*_*nn*,∞_ ≡ *α_nn_*(*t* → ∞).

We lack the data to fit exactly the plateau value, and we set the reasonable value *α*_*nn*,∞_ = 0.8. Least-square error estimation of the two parameters yields: *τ_nn_* = 79.3 hpf and *σ_nn_* = 14.5 hpf (sq. error = 0.03).

The pn-divisions are constrained to be complementary to 1 of the two others:

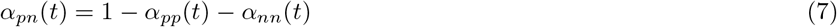

and its square error is 0.02.

We report the corresponding evolution of *γ*(*t*) in Fig. 1-b.

With these profiles for *α_pp_*(*t*) and *α_nn_*(*t*), the evolution of the P-pool evolves according to (details in Methods Eq 15):

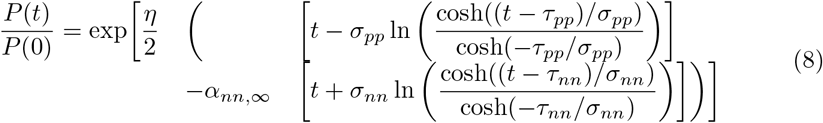

The dynamics of the progenitors and neurons pools, driven by this balance, also depends on the initial condition and the division rate. Setting *P*(0) = 1, *N*(0) = 0 at time *t*_0_ = 44 hpf and *η* = 1/12 hours [13, 9], this system yields a good account of the evolution measured by Saade et al.[12] (Fig. 1-c). At the beginning, the large bias towards proliferative divisions amplifies the pool of progenitors up to a maximum value : *P_max_*[*CTL*] = 5 per initial progenitor at around *t_maxp_*[*CTL*] = 72 hpf, corresponding to the change of sign of *γ*(*t*). Then, the production of neurons raises mainly due to pn-divisions until nn-divisions become dominant over pn-divisions (at around 82-83 hpf). The pool of progenitors depletes to zero while terminal divisions increase the pool of neurons up to a plateau value of *N*(*t* → ∞)[*CTL*] = 17.6 neurons per progenitor initially present. We note that this evolution, and especially *N*(*t* → ∞) is highly sensitive to the value of *t*_0_.

#### 2.1.1 Incorporating CDC25B experiments

Bonnet et al.[9] have performed a series of experimental manipulation of the expression of CDC25B phosphatase in this biological system. Their experimental measures are the proportions of progenitors / neurons, and a corresponding measure of the modes of division, depending on the experimental condition : Control (CTL), Gain of Function (CDC25B GoF) using the wild-type form of CDC25B, Gain of Function using a CDC25B modified not to be able to interact with known substrates CDKs (CDC25B^ΔCDK^ GoF).

Modes of division were measured by Bonnet et al.[9] at stage HH17, and fit well with the MoDs measured by Saade et al.[12] at time 72 hpf (Fig 1-a, circle dots). To make the correspondence between P/N fractions reported in Bonnet et al.[9] and the P/N evolution measured in Saade et al.[12], we had however to consider that the former correspond respectively to times 60 hpf and 84 hpf on the time scale in Saade et al.[12] (i.e. 12 h before and after 72 hpf, keeping the correct interval of 24h in between).

To check the power of this simple model, we now explore the hypothesis that CDC25B GoF has only an effect upon the schedule of MoD transitions. We expect that GoF would shift the balance *γ*(*t*) sooner in time, and indeed, the measured MoD in the GoF experiment can be fitted by shifting the three time profiles 8 hours sooner (Fig 2-a).

**Figure 2.**
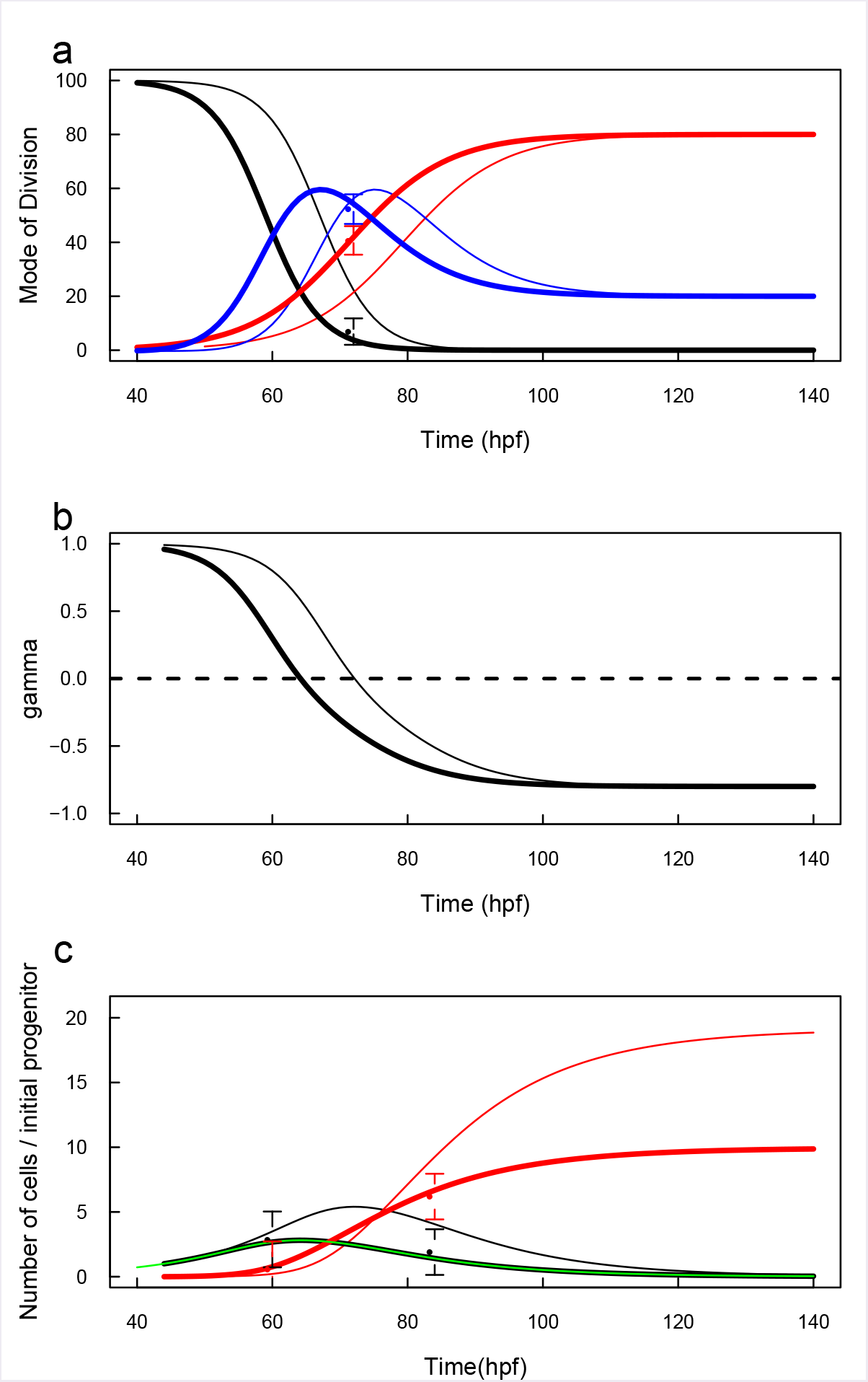
CDC25B Gain-of-Function promotes neurogenic divisions. Bonnet et al. measured MoD in presence of CDC25B GoF (dots and CI). a) In a minimalistic approach, we considered that GoF only hasten the transition from proliferation to differentiation, so that the PP and NN profiles were shifted to fit the measured GoF MoD (thick lines) 8 hours sooner than the CTL profiles (thin lines). PN profile was adjusted accordingly, b) Corresponding evolution of the balance proliferation / differentiation γ (GoF: thick line, CTL: thin line). c) Predicted evolution of the pools of progenitors (black) and neurons (red) under GoF (thick lines) compared to CTL (thin lines). The dots report the proportion of progenitors / neurons measured in Bonnet et al. in GoF condition, scaled to the total amount predicted at their respective times. We note that GoF promotion of neurogenic divisions results in a large decrease of the final pool of neurons, because progenitors lack time to amplify.

Interestingly, at time 72 hpf, this strongly affects *α_pp_* and *α_nn_* but leaves *α_pn_* unchanged. The corresponding evolutions of the pools P/N are strongly affected, since the progenitors lack time to proliferate, reaching now a maximum of *P_max_*[*GoF*] = 2.6 per initial progenitor at around *t_maxp_*[*GoF*] = 64 hpf. As a consequence, the pool of neurons increases sooner, but reach a plateau value nearly twice as less as in the CTL condition, *N*(*t* → ∞)[*GoF*] = 9.2 neurons per initial progenitor. The proportions P/N measured by Bonnet et al.[9] fit well with this picture.

The case of CDC25B^ΔCDK^ GoF yields a different picture. Here, the pp-divisions had to be advanced by 2 hpf while the nn-divisions had to be delayed by 4 hours to correspond to the ones measured by Bonnet et al.[9] (fig. 3a). As a result, the main effect of CDC25B^ΔCDK^ GoF is to greatly promote pn-divisions, and they appear sooner and reach a higher proportion. This suggests that CDC25B^ΔCDK^ GoF promotes asymmetric neurogenic pn-divisions, but fails to promote the transition from pn-divisions to nn-divisions as does CDC25B GoF. Remarkably, since the pn-divisions are neutral to the balance proliferation / differentiation, this poorly affects its evolution (fig. 3b), which in turn translates to almost identical evolution of the P/N pools (fig. 3c). Here again, the proportions P/N measured by Bonnet et al.[9] fit well with this picture. We note that the effect of CDC25B-ΔCDK could not be detected by measuring only the P/N pools evolution.

**Figure 3.**
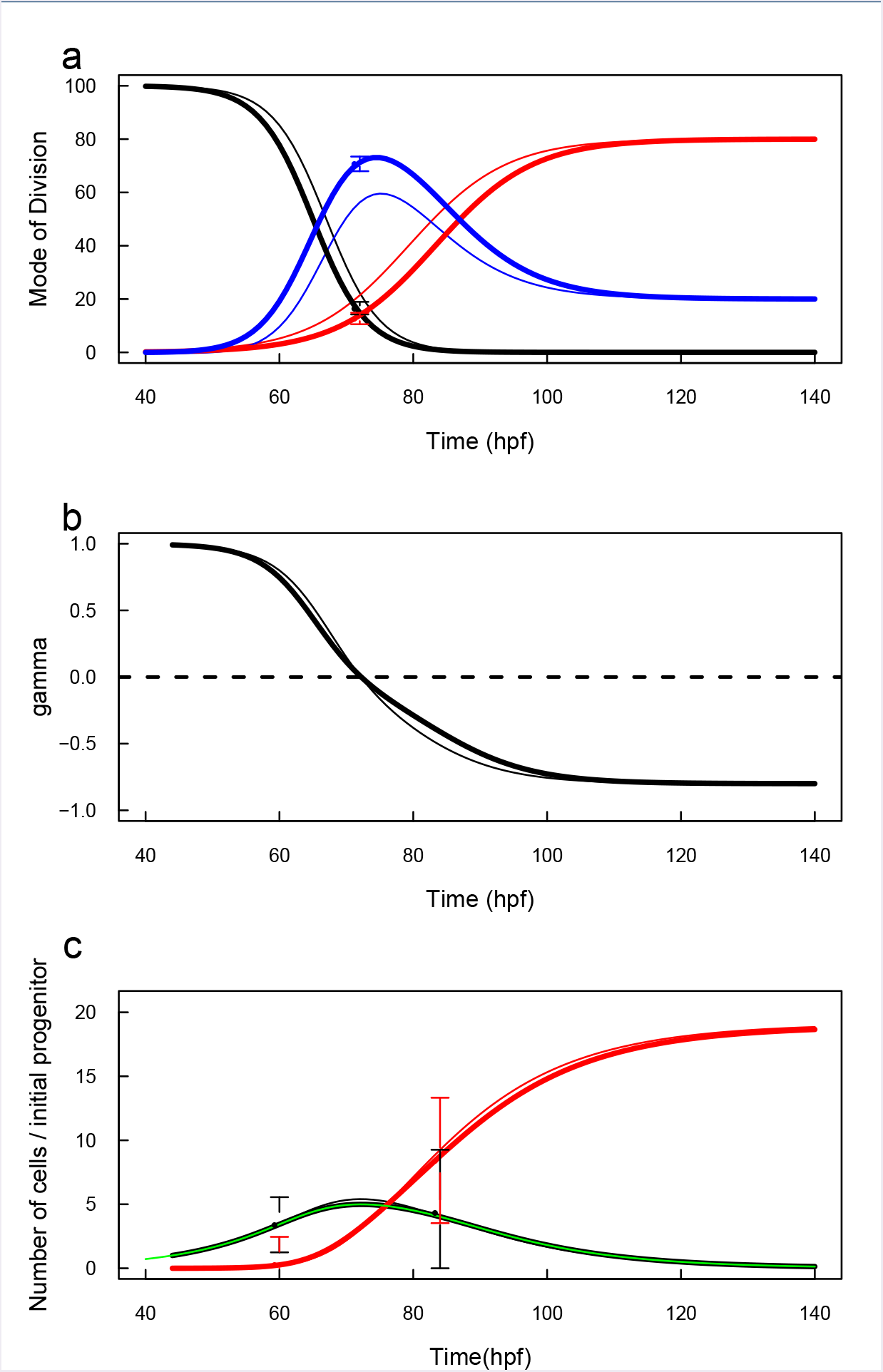
CDC25B-Δ CDK Gain-of-Function have a differential effect upon neurogenic divisions. a) Bonnet et al. measured MoD in presence of CDC25B-Δ CDK GoF (dots and CI), suppressing the interaction between CDC25B and its cyclin. To fit the measured data by only shifting profiles in time, the PP profile has to be shifted sooner (-2 hours) and the NN profile has to be shifted later (+4 hours). As a consequence, the complementary PN profile is enhanced (compared to the CTL) and lasts longer. b) The balance proliferation / differentiation is poorly affected (because PN-divisions are neutral for the balance). c) The dynamics of the two pools is very close to the CTL dynamics and match with the measured proportions given in Bonnet et al.

Together, the model given by the system 1 expresses the dynamics at the population scale, yielding the evolution of the two kinds of cells: the pool of progenitors, and the pool of neurons. Being formulated at the population scale, the variables and the parameters represent averages over large ensemble of cells (or over repeated replications). At the cell level in the biological system, those averages can correspond to numerous scenarios. Nonetheless, the model dynamics produced by Eqs. 5, 6, 7 should be taken as a point of reference because any scenario at the cell scale should reproduce these dynamics at the population scale. In that sense, the basic model should be regarded as a way to describe a strong constraint over the set of possible cell-scale scenarios and a guide to narrow the research of mechanistic explanations. In the next section, we will use it as such in order to explore three scenarios incorporating fate restriction at the cell scale.

### 2.2 Models with fate restriction

The basic model 1 is compatible with the simplest interpretation at the cell level: that each dividing cell is liable to stochastically produce the three possible MoD, in proportion to what is measured at the population scale. We now explore alternative models in which we introduce fate restriction. Fate restriction denotes the fact that some dividing cells cannot produce all possible MoD anymore. To incorporate fate restriction in model, we have then to consider that the pool of progenitors is actually composed of different kinds of dividing cells, each kind being able to produce only a restricted set of MoD.

Let consider the case with only two sub-populations of dividing cells. In this case, restricting sets of MoD entails that one of the population cannot perform purely self-replicating division anymore, and must then be restricted to a choice between asymmetrical neurogenic (self-renewing) or terminal (self-consuming) MoD. We denote this sub-population *A*(*t*), and we denote *G*(*t*) the other one, with *G*(*t*)+*A*(*t*) = *P*(*t*), where *P*(*t*) is the total pool of dividing cells (progenitors in the model 1). In short, fate restriction translates in the constraint that a daughter of a cell of the pool *G* and that becomes of type *A* cannot reverse its fate to become back of *G*-type. We note that, at this stage, the two pools *G*(*t*) and *A*(*t*) are distinguished from a model structure standpoint, not a biological one.

The possible scenarios we see are:

1. GAA-model: *G* → (*G, G*), *G* → (*A, A*), *A* → (*A, N*), *A* → (*N, N*) *G*-cells are capable of self-duplication or symmetrical non-neurogenic division. During the latter MoD, the two daughter cells become of type A, with fate restricted to asymmetrical self-renewing or terminal differentiation.
2. GAG-model: *G* → (*G, G*), *G* → (*A, G*), *A* → (*A, N*), *A* → (*N, N*) *G*-cells are capable of self-duplication or asymmetrical non-neurogenic division. During the latter MoD, one cell becomes of type A, with fate restricted to asymmetrical self-renewing or terminal differentiation.
3. GAN-model: *G* → (*G, G*), *G* → (*A, N*), *A* → (*A, N*), *A* → (*N, N*) *G*-cells are capable of self-duplication or asymmetrical neurogenic division. A dividing cell issued from the latter becomes of type A as it cannot self-duplicate anymore and is restricted to asymmetrical self-renewing or terminal differentiation.

The only difference between the three scenarios is the non self-replicating MoD of the *G*-cells (hence their names). The difference could involve some mechanisms acting during this MoD. In scenario GAA, the symmetrical output of the MoD suggests some kind of time-related events, such as external signaling or internal generation markers. In scenario GAG, some stochasticity during the cell cycle could be instrumental to induce the loss of self-replication power in one daughter cell, such as a lack of perfect replication. In scenario GAN, the neurogenic mechanism producing the N-cell would be instrumental to induce fate restriction in the companion daughter A-cell.

We examine below the three scenarios in the light of the data presented above. We use the evolutions of MoD in the PN model to calibrate the MoD in each scenario for the three conditions (CTL, CDC25B GoF, CDC25B^ΔCDK^ GoF), and check how it accounts for the evolution of P/N cells. In the end, we examine how the three scenarios are structurally compatible with the observed evolutions reported by Saade et al.[12].

#### 2.2.1 Model GAA

This model obeys:

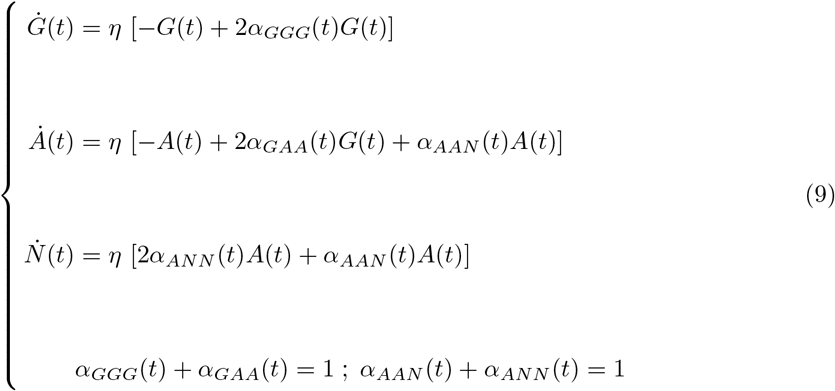

and the correspondences between GAA scenario variables and the variables in model 1 are:

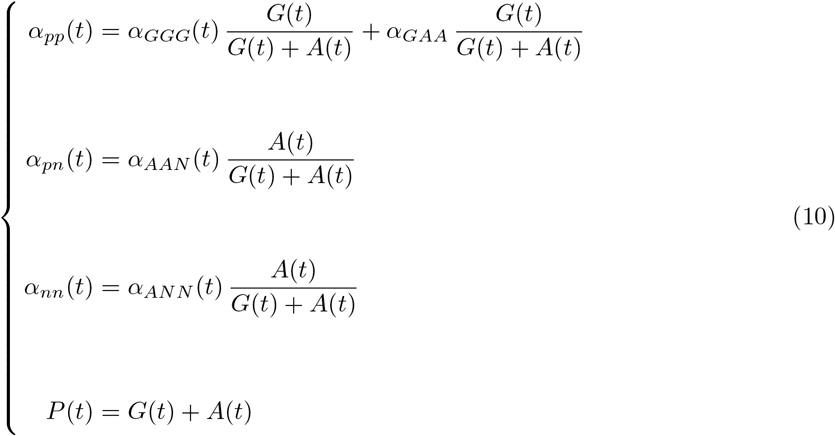

To establish this correspondence, we have considered that the observed proliferative MoD *P* → (*P, P*) in model 1 aggregates all underlying proliferative divisions by the *G* pool : *G* → (*G, G*) and *A* → (*A, A*). The observable *α*_.._ (*t*) functions express the proportions of each MoD among a total number of divisions. They can be regarded as a probability that a given division is of a given kind of MoD. Hence, to reconstruct a given observable MoD, we have to combine the probability the corresponding kind of progenitor would adopt this MoD with the proportion of this kind of progenitors among the total number of progenitors. For instance, the probability observing a PN division, *α_pn_*(*t*) (the observable proportion of asymmetric neurogenic divisions), is the probability that a given progenitor is of type *A* (namely *A*(*t*)/(*G*(*t*) + *A*(*t*))) times the probability that this progenitor performs an AAN division (*α_AAN_*(*t*)). We proceed this way for the three kinds of observable MoD.

We used MoD fitted in model 1 to calibrate the four MoD functions *α_GGG_*(*t*), *α_GAG_*(*t*), *α_AAN_*(*t*) and *α_ANN_*(*t*). The procedure is given in full details in Methods. The resulting evolutions are given in Fig. 4 (please note that for programming facility, we shifted the time scale so that 44 hpf corresponds now to time 0).

**Figure 4.**
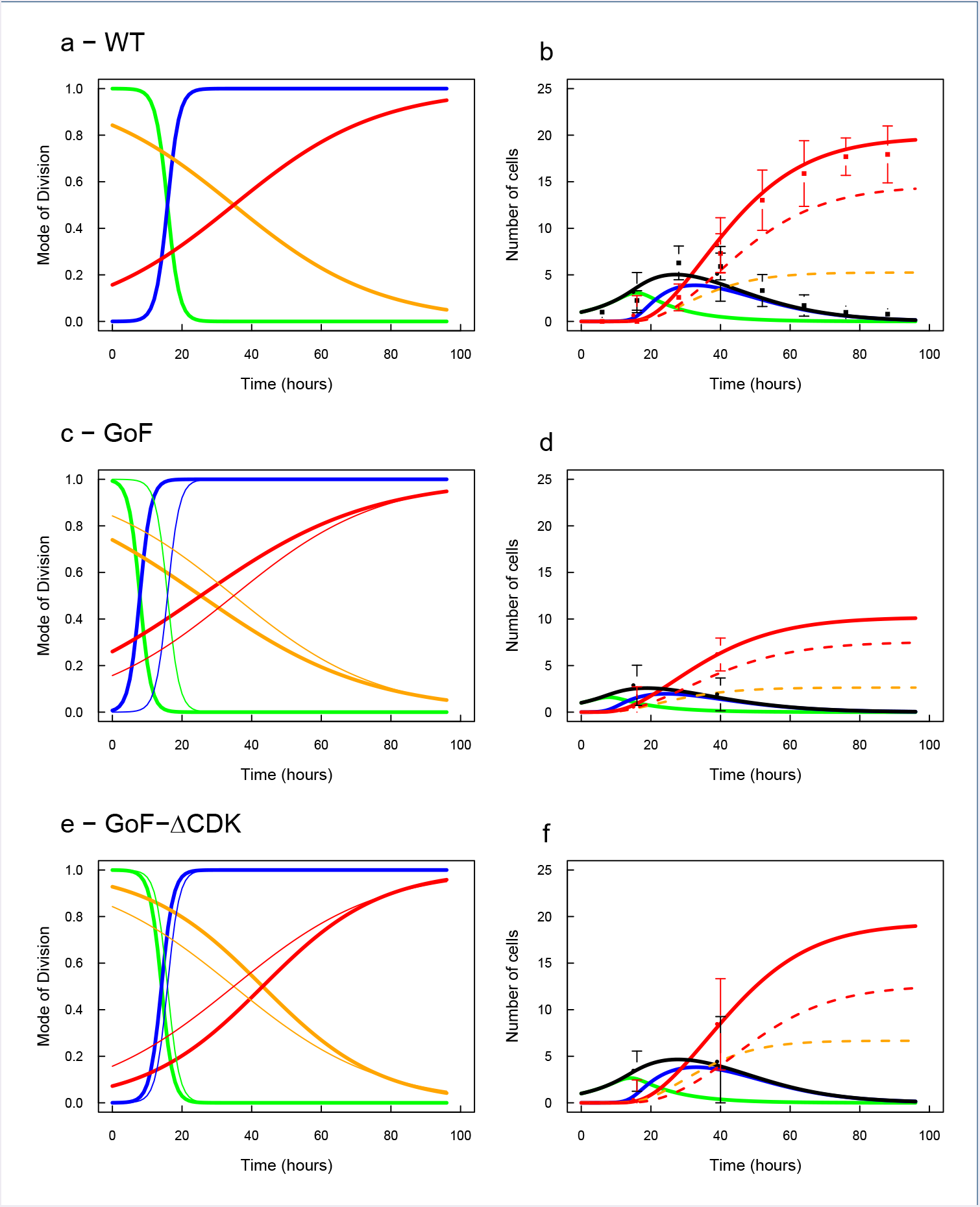
Model GAA. Left column: evolution of the GAA-MoD constrained by the MoD data (green:*G* → (*G, G*), blue:*G* → (*A, A*), orange:*A* → (*A, N*), red:*A* → (*N, N*)). Right column: predicted evolutions for the three cell pools (green:*G*(*t*), blue:*A*(*t*), black:*G*(*t*) + *A*(*t*), red:*N*(*t*), points: data). Origin of time corresponds to 44 hpf. The thin lines in c and e are the profiles from a, for eye-comparison. Dotted lines on right column indicate origin of neurons: orange dotted lines for neurons issued from *A* → (*A, N*) MoD, and red dotted lines for neurons issued from *A* → (*N, N*) MoD.

Under CTL condition, we observe an abrupt and early switch of the *G*-cells MoD, from dominant *G* → (*G, G*) MoD (Fig. 4a, green) before 16h to dominant *G* → (*A, A*) MoD (Fig. 4a, blue) after 16h (that would correspond to 16 + 44 = 60hpf). As a consequence, the P-pool is made of only *G*-cells up to that time (Fig. 4b, black and green curves), and there is a clear switch after which *A*-cells become dominant in the system (Fig. 4b, blue curve). Contrastingly, the MoD of *A*-cells evolves smoothly (Fig. 4a, orange and red) and the characteristic time of their switch (the moment when the two curves cross each other) is about 35h (79hpf). This leave time for A-cells to produce neurons by *A* → (*A, N*) MoD (Fig. 4b, dotted orange) so that neurons issued from asymmetric neurogenic divisions will represent 25% of the N-pool in the end. After 35h, *A*-cells engage in terminal differentiation up to extinction.

The evolutions of *P* = *G* + *A* and *N* pools produced by these calibrated MoD match very well the measured ones.

In CDC25B GoF condition, the 8-hours advance of MoD in model 1 is directly reflected in the MoD for the *G*-cells (Fig. 4c). This is expected given the calibration procedure, and this is true also for the switching time of the MoD for the *A*-cell, although their slopes are further smoothened. This results into P/N profiles under GoF condition that reflect the profiles under model 1 (Fig. 4d).

In CDC25B^ΔCDK^ GoF condition, the switch of MoD for the *G*-cells happens slightly sooner than in CTL condition (Fig. 4d, green and blue), so that the total number of A cells produced by *G* → (*A, A*) is a bit lower (and hence for P = *G* + *A*-cells). On the contrary, the MoD for the *A*-cells are delayed by about 5 hours (Fig. 4e, orange and red). This is consistent with the observation than pn-divisions in model 1 are favored under CDC25B^ΔCDK^ GoF condition where they operate for a longer time than in the CTL condition. This results into a larger production of neurons through asymmetric neurogenic divisions *A* → (*A, N*) (Fig. 4f, dotted orange). By contrast, since the total number of *A*-cells is lower than in CTL condition, so is the number of neurons produced by *A* → (*N, N*) MoD (this number is logically twice the total number of *A*-cells produced in the process). Eventually, both compensate and the total number of neurons is in the end the same as in CTL condition.

Overall, the structure of this model appears compatible with the data. It is characterized by an early switch in the MoD of *G*-cells, and a late and smooth transition in the MoD of *A*-cells. The effect of CDC25B GoF would be to accelerate the transition of MoD for both *G* and *A* cells by the same amount, and the effect of CDC25B^ΔCDK^ GoF condition would be to delay only the MoD of *A* cells, leaving MoD of *G* little affected.

#### 2.2.2 Model GAG

This model obeys:

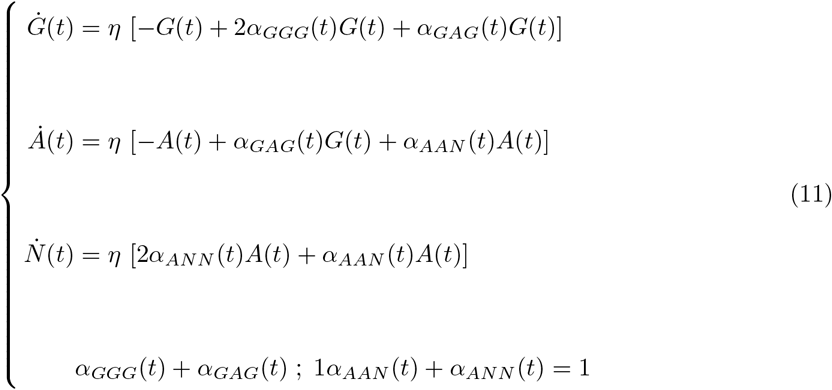

The correspondence between scenario variables and the variables in model 1 is given by:

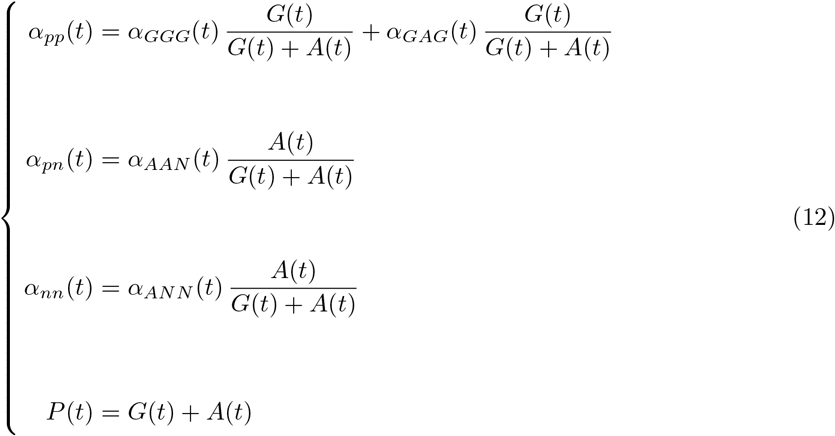

where we have considered that the observed proliferative divisions *P* → (*P, P*) in model 1 aggregate all proliferative divisions by the *G* pool : *G* → (*G, G*) and *A* → (*A, G*). We used the same procedure to calibrate the four unknown functions *α_GGG_*(*t*), *α_GAG_*(*t*), *α_AAN_* (*t*) and *α_ANN_* (*t*) from MoD fitted in model 1. Full details are given in Methods. The resulting evolutions are given in Fig. 5.

**Figure 5.**
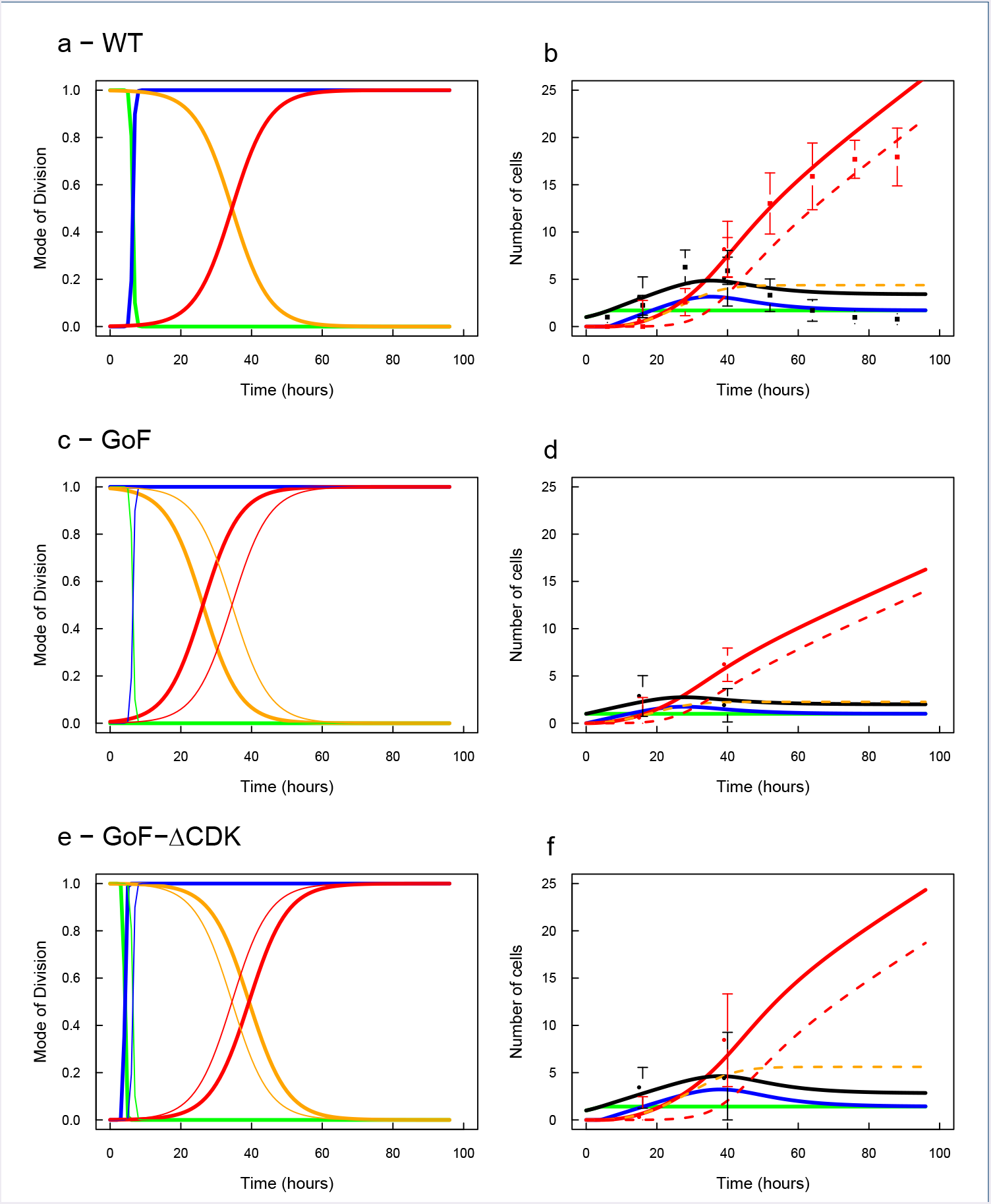
Model GAG. Same legend as Fig. 4, except that blue in the left column is *G* → (*A, G*).

The calibration procedure for the MoD of *G*-cells in the CTL case suggests an almost perfect switch at time 6.4h (50.4hpf) and here again a smoother evolution for the MoD of *A*-cells (Fig. 5a), in the same line as model GAA. However, observing the corresponding evolution of *P* = *G* + *A* and N pools (Fig. 5b), it is obvious that this model structurally misbehaves. Indeed, the switch in *G*-cells MoD triggers *G* → (*A, G*) MoD which are self-renewing for ever, so that the *G*-pool cannot decrease anymore. This MoD also produces *A*-cells, and since it does from a stabilized *G* pool, it continuously produces *A*-cells. Before the switching time of the *A*-cells MoD, the newly produced A cells produce neurons through asymmetric neurogenic divisions *A* → (*A, N*) (Fig. 5b, dotted orange), but after that time, all newly produced A cells differentiate into 2 neurons. Hence, the dynamics become trapped in a perpetual regime where *G*-cells are continuously producing *A*-cells, which in turn continuously contribute to an increase of the *N* pool (Fig. 5b, dotted red). We note that the structural default of GAG model becomes inconsistent with measures in CTL only after time 60h (104 hpf).

In CDC25B^ΔCDK^ GoF condition, this structural inadequacy translates into an aberrant fit for *G*-cells MoD (fitted switching time being-613h). This is not surprising given the diagnostic above that the procedure tries to fit a monotonously increasing function to the increasing-decreasing evolution of the *P* pool predicted by the MoD fitted in PN model. Over the considered period of time, this means that the best value for *G*-MoD are just an absence of *G* → (*G, G*) MoD, as if the system would adopt the perpetual regime from the start. We note however that even under this badly conditioned model, the effect of CDC25B GoF would translate into an advanced transition time for the MoD of *A*-cells, and that the early evolution of *P* = *G* + *A* and *N* cells are compatible with data.

The same is true for CDC25B^ΔCDK^ GoF condition where the calibration suggests that this condition poorly affect the MoD of *G*-cells, and delays the MoD of *A*-cells and yields evolution of *P* = *G* + *A* and *N* cells that are compatible with data measured in the early phase.

Overall, this model is just a good indication that our calibration procedure does not allow *any* model to fit. Despite its structural default could have been stated directly from its evolution equations, we consider it was worth mentioning it because, if experimental data were pointing to it, it would call for additional mechanisms to control and stop the perpetual regime.

#### 2.2.3 Model GAN

This model obeys:

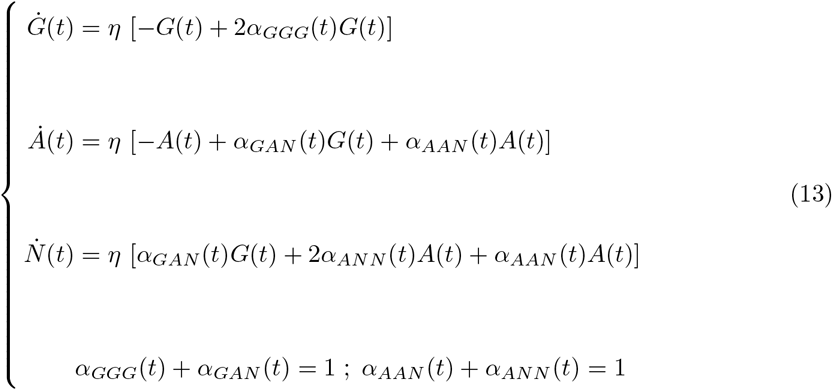

The correspondence between scenario variables and the variables in model 1 is given by:

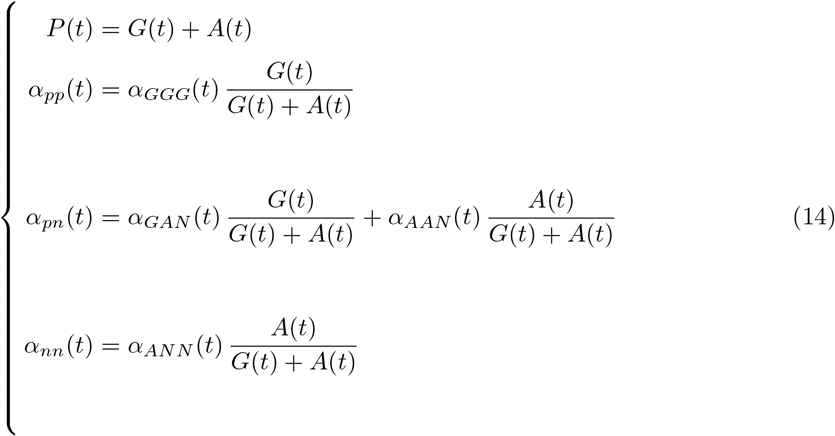

where we have considered that the observed asymmetric divisions *α_pn_* aggregate the asymmetric divisions *G* → (*A, N*) by the *G* pool and asymmetric divisions *A* → (*A, N*) by the *A* pool.

We used the same procedure to calibrate the four unknown functions *α_GGG_* (*t*), *α_GAG_*(*t*), *α_AAN_* (*t*) and *α_ANN_* (*t*) from MoD fitted in model 1. Full details are given in Methods. We note that due to the one-to-one correspondence between *α_pp_* and *α_GGG_* in this scenario, the *α_GGG_* (*t*) profile could be analytically retrieved by an analytical inversion. We then checked that the least-square error procedure to calibrate *α_GGG_*(*t*) used for the two models above yielded the same result.

The resulting evolutions are given in Fig. 6.

**Figure 6.**
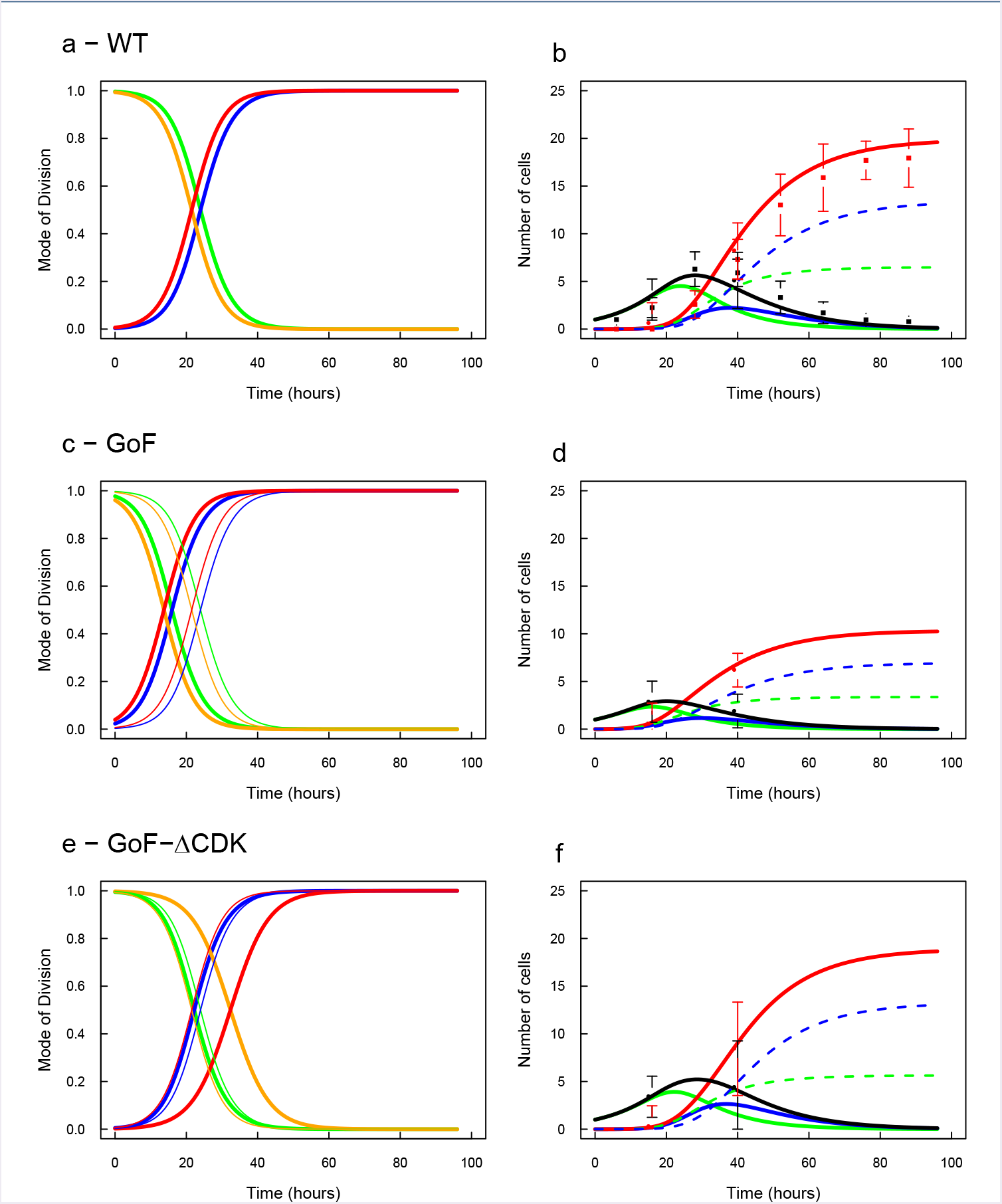
Model GAN. Same legend as Fig. 4, except that blue in the left column is *G* → (*A,N*) Dotted lines on right column indicate origin of neurons: green dotted lines for neurons issued from *G* → (*A,N*) MoD, and blue dotted lines for neurons issued from *A* → (*A,N*) MoD.

In the CTL case, we found a remarkable convergence of the MoD evolutions for *G*-cells and *A*-cells and we recover a perfect prediction for the evolution of *P*(*t*) and *N*(*t*) populations.

The typical switching times of MoD is 24h for the *G*-cells and 21.5h for the *A*-cells (i.e. 68hpf and 65.5hpf), and their switching rates are practically identical. In the beginning, the G-pool is mainly proliferating, while *G* → (*G, G*) is dominant over *G* → (*A, N*), for about 20 hours (Fig. 6a, green vs. blue). This yields a growth of the *G* pool up to a peak at 4.5 *G*-cells (per initial *G*-cells) at 24 hours (Fig. 6b, green). They represent 88% of *P*-cells at that time.

After that peak, *G*-cells slowly decreases while populating *A* and *N* cells through *G* → (*A, N*) divisions. From that time, *A*-cells are produced, but their MoD are already very skewed in favor of *A* → (*N, N*) (Fig. 6a, red vs. orange) so that they are almost consumed by terminal divisions as soon as they are produced.

Seeing this, we checked a simpler scenario *G* → (*G, G*), *G* → (*A, N*), *A* → (*N, N*) so a progenitor issued from an asymmetrical division would always differentiate in two neurons at the next cycle. This yields practically the same results (not shown).

In the CDC25B GoF case, the 8-hours advanced evolution of the MoD in model 1 directly translates into an equivalent and parallel 8-hours advanced evolution for *G* and *A* MoD, which is not surprising given the estimation method.

Contrastingly, the evolution of these MoD differs in the case of CDC25B^ΔCDK^ GoF. As expected, the small advanced *α_PP_* profile affects little the switching time of the *G*-cells. However, the 4-hours delayed *α_NN_* profile translates into a threefold larger delay for the *A*-cells MoD, namely they are shifted 11-hours later than in the CTL condition (32.4h vs 21.5h). As a consequence, the *A* → (*A, N*) MoD becomes operative since it is still around 0.5 when *A*-cells reach their peak. In the end, the production of neurons is slightly increased by 18.8% of the CTL value.

Overall, this model clearly appears as the simplest one to explain. The MoD of *G* and *A* cells would evolve at the same pace in the CTL condition (hence possibly under the control of the same signal). CDC25B GoF would accelerate them the same way. CDC25B^ΔCDK^ GoF would only delay MoD of *A*-cells.

## 3 Discussion

Our question was to test whether the neurogenic progenitors observed in the developing spinal cord (should they do an asymmetric or a terminal division) could correspond to a particular set of cells that would be characterized by their loss of proliferative power (fate restriction).

To this end, we have first established a general restriction-free model with progenitors able to perform any division (model 1). Fitting the evolution of its MoDs (PP, PN, NN) from data published by Saade et al [12], we found smooth MoDs time-profiles that can account for the evolution of the P and N pools reported in [12].

We characterized the behavior of this model under manipulative experiments made by us [9] with CDC25B and this gives support to the hypothesis that the action of this phosphatase is reflected by delayed or advanced MoD switching times. We consider this general model as a reflect of the observables that are presently available using Sox2 / Tis21 and HuCD staining. We take it as a benchmark to constrain refined scenarios with heterogeneous progenitors. We note that its general structure is also compatible with a broad description of progenitors / neurons evolution in the neocortex [14, 15]. It should hold as well for other neural tube zones, such as the dorsal area where CDC25B is expressed at the peak of neural production [16, 9]

Next we have explored three model structures embedding a fate restriction in progenitors, introducing two different progenitors population with the structural constraint that one of them cannot do self-replication. For the three models compatible with this constraint, we have derived the corresponding system of evolution equations. We have established the correspondence between the evolution of each restricted but not observable MoDs and the evolution of the unrestricted but observable MoDs of the benchmark model 1. This correspondence was used to calibrate the evolution of the restricted MoDs as close as possible to the observable MoDs, and we checked the resulting behavior of the model regarding the evolution of observable P/N cells pools. Two models, GAA and GAN, appear structurally compatible with the P/N data and yield acceptable predictions.

To refine further model diagnosis, we now look back at how close the hidden MoD (and balance) could be actually fitted to the observable ones (Fig. 7). A visual inspection is sufficient to give preference to the GAN model, even if GAA model can still not be formally excluded, especially regarding the balance proliferation / differentiation.

**Figure 7.**
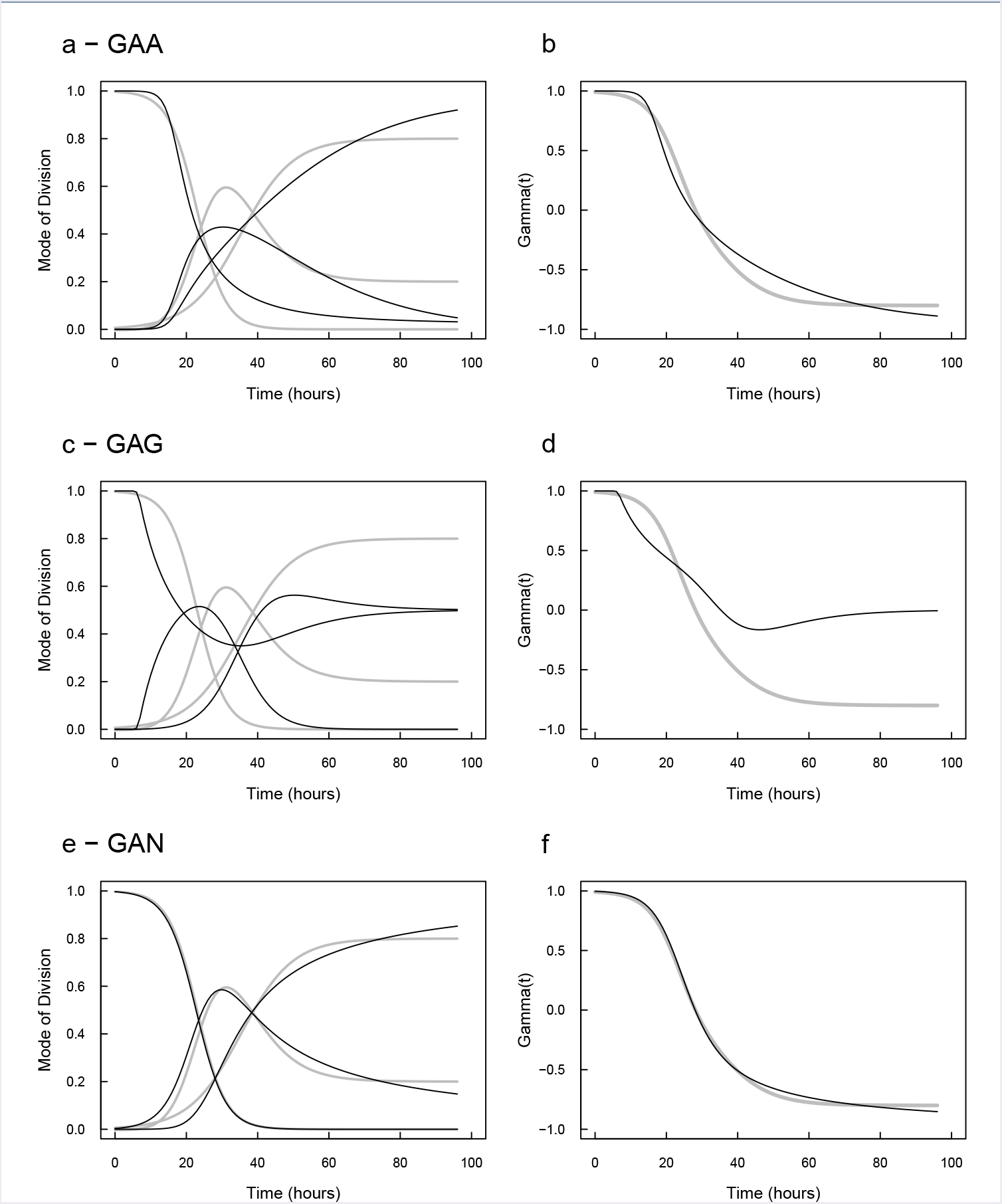
Structural compatibility of each model regarding the MoDs and proliferation/differentiation balance. (left) MoDs reconstructed under each model (black lines) are to be compared with the observable MoDs. (right) Corresponding evolution of the balance proliferation / differentiation. If the model GAA can not be formally excluded (especially regarding the resulting balance), the GAN model appears as the most compatible with the observed MoDs.

Ultimately, we illustrate how fate-restricted GAN model could perform better than the unrestricted model 1 (Fig. 8). They cannot be distinguished by their predictions regarding the evolution of P/N pools (Fig. 8c,d, the P-pool for GAN being *P* = *G* + *A*), neither by the evolution of PP-divisions which are very close to each other. We see however a difference in PN and NN MoDs at the beginning of the process where NN divisions rise up later in GAN model than in model 1 and seem more adequate. This difference is due to the fact that NN divisions cannot appear before the A-pool has increased, whereas they can happen earlier in model 1 through NN divisions. This effect yields in the end a better fit of the MoDs under GAN model. Importantly, we note that this better fit is not due to differences in degrees of freedom (free parameters) for the MoD, since both models have four, so it is attributable only to their different structures : in GAN model, the PN divisions are GAN divisions convoluted by the population of *A*-cells.

**Figure 8.**
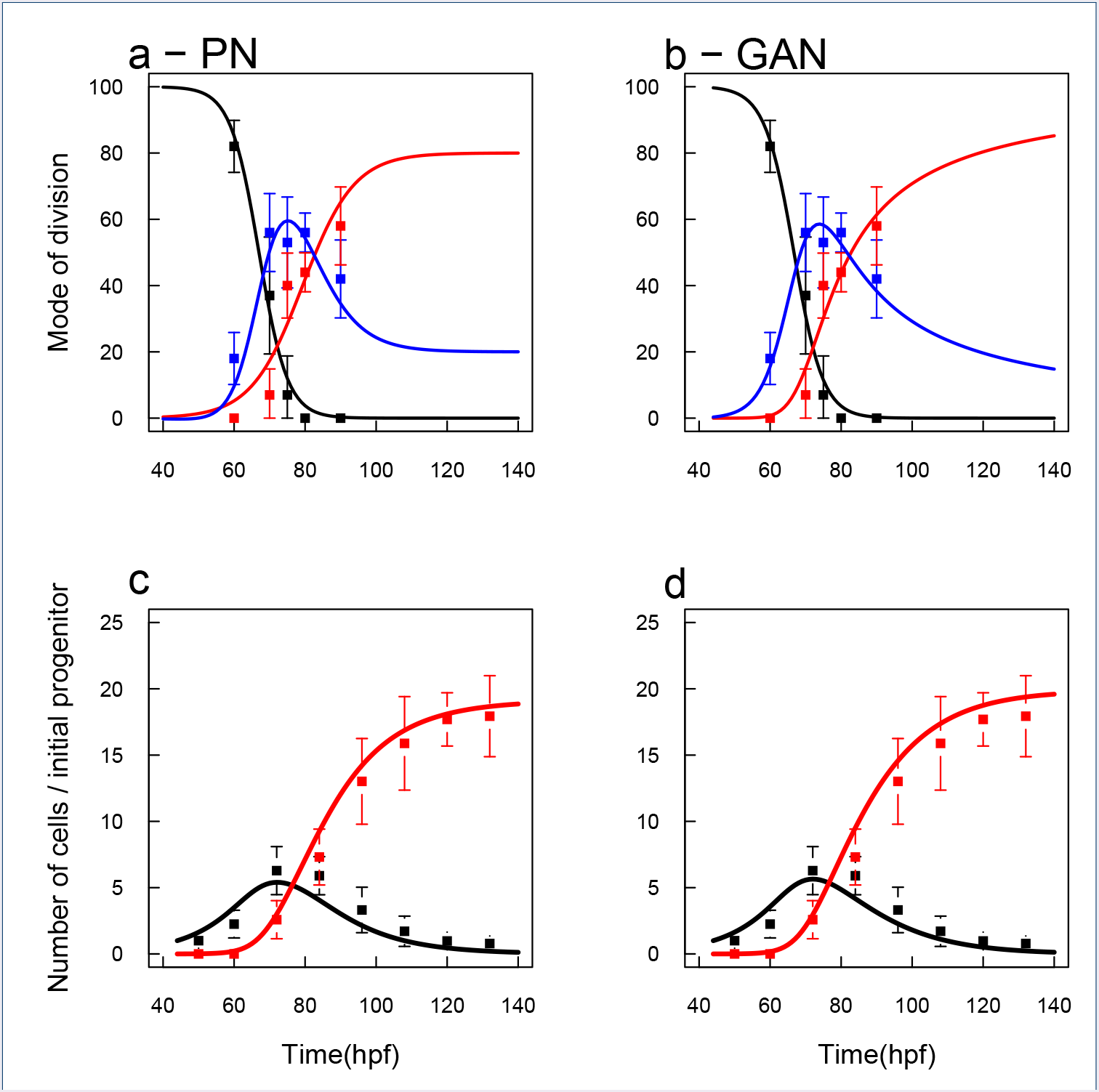
Models PN and GAN vs data. (a) and (b) Observable PP/PN/PP MoDs for the two models. (c) and (d) Observable P/N populations for the two models. In (a), for PN model, are represented directly the fitted MoDs. In (b), for GAN model, are represented the MoD reconstructed from Eq.14, and which result from the combined effect of *G / A* dynamics and their respective MoDs. (b) vs (a) suggests than GAN model behaves better than PN model, especially in the beginning of the process (ca 60hpf) because symmetric neurogenic MoD (red) has to wait for *A*-cells to be produced. (d) vs (c) shows that both models yield the same correct predictions.

From a modeling standpoint (where modeling is used as a way to gain clarity in the face of intricacy), GAN model displays several interesting features compared to model 1. First, it can be considered as simpler to be mechanistically explained since both types of progenitors display the same evolution of their MoDs (with the same transition time and same slope), and this evolution is monotonous. In contrast, model 1 would call for a specific explanation of the non monotonous evolution of PN divisions as well as explanation of the complicated transitions among MoDs. Secondly, CDC25B GoF effect is the same for both models: it makes transition times happen earlier, and with the same extent. Thirdly, CDC25B^ΔCDK^ GoF effect is interpreted straightforwardly in GAN model : the phosphatase unable to interact with its CDK substrate just delays the transition time of *A*-cells (it maintains A-cells in self-renewing mode for a longer time). In model 1, CDC25B^ΔCDK^ GoF effect appears as compound and would ask for a complicated explanation for the differential effect upon advanced PP and delayed NN divisions.

We note that our modeling proposition displays an important difference with the model proposed by Saade et al. themselves [12] (see also [17]): we do not detect a strong switch of MoD at the population level. Their basic model incorporates an all-or-nothing switch at time *t*\* ≃ 80 hpf with only proliferative divisions (PP) before *t*\* and only neurogenic divisions (PN or NN) after *t*\*. This is equivalent to a fate restriction that would apply to progenitors, all at once, at time *t*\* (in terms of GAN model, all *G*-cells would become *A*-cells at time *t*\*). They next extend this model to allow smoother transitions, division asynchrony, accelerating cell cycle and a *de novo* incorporation of new progenitors under the induction of Shh. Even with this smoother model, their fitting yields a sharp extinction of PP-divisions at 73 hpf (from 60% to 0% within one hour). It is difficult to determine how this finding is constrained by the initial choice in their basic model, but this predicted evolution of the MoD appears at odd with the observed ones and can predict a sensible evolution of the P/N populations only due to the additional source that compensates for the sharp extinction of proliferative divisions.

We observe that our model does not incorporate a source of progenitors so the structures of the models are different. We also note that the fitting procedures were not the same. Saade et al. fit the 13 free parameters of their extended model using *an error minimization algorithm with respect to the experimental data* [12] (Extended Experimental Procedures — Mathematical Modeling). As we understand this sentence, they fit the MoD profiles and the source intensity so that the predicted dynamics of the P/N populations matched as close as possible the observed evolution. We have proceeded differently: we have minimized the error between the modeled MoD evolution and the observed MoD evolution, and then only have we checked how the predicted P/N evolutions match or not the observed ones. As a consequence of our procedure, the MoD profiles in the PN and in the GAN models are by construction as close as possible of the observed MoD. Importantly, both procedures have to set an initial condition (i.e. an absolute time 0 at which we fix the initial pool of progenitors), and since proliferative processes are exponential, evolutions of P/N populations are highly sensitive to that choice. We guess that a small change (by more or less two hours) of that time 0 in Saade et al. model would have a strong effect to the required intensity of their additional source. Conversely, our model being in the same extent sensitive to the choice of time 0, a different choice (by more or less two hours) could be compensated for by an additional source if needed. The crucial point here is that the relative error for experimental data is the highest at that early time, because there are few progenitors then, and the developmental stage is only determinable with an error of the same extent (more or less two hours).

Finally, we advocate that our model indeed incorporates a switch mechanism, but it is specified at the cell level: the switch operates for a given G-cell when it becomes an *A*-cell. Under the hypothesis of asynchronous divisions, the smooth evolution of MoD profiles reported for the GAN model is then compatible with an abrupt signaling event: cells that have undergone a *G* → *A* switch after this signaling event would only display their new division modes (AAN, ANN) at their next M-phase. At the population scale, the switching between MoD would then require at least one cell cycle time length to fully display. This is about the order we observe in the GAN model where the MoD transition happens over about 20 hours, i.e. one to two cell cycle lengths (12 hours).

## 4 Methods

### 4.1 Solving *P*(*t*) from Eq.4

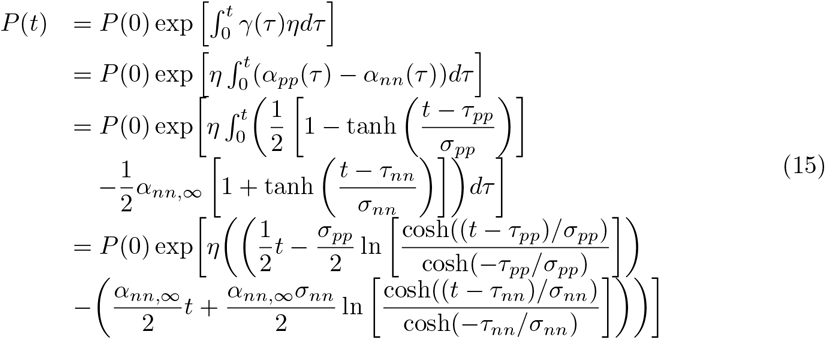

hence :

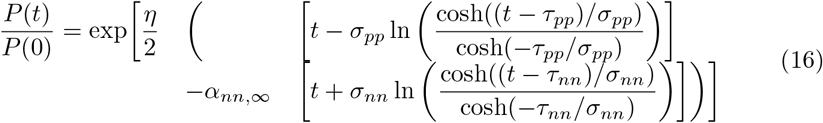

### 4.2 GAA inversion

#### 4.2.1 Estimating *α_GGG_*(*t*)

In principle, we should have:

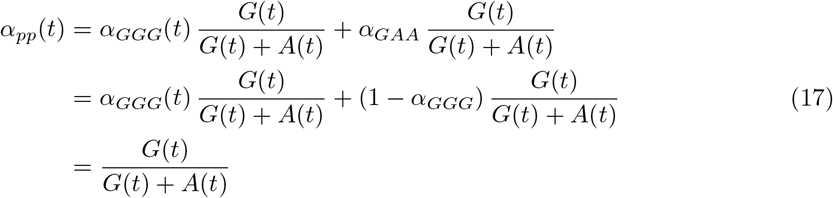

where we have used that *α_GGG_*(*t*) + *α_GAA_* = 1.

Using *P*(*t*) = *G*(*t*) + *A*(*t*), we obtain:

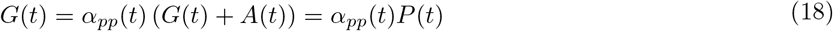

Since

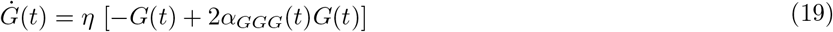

we have:

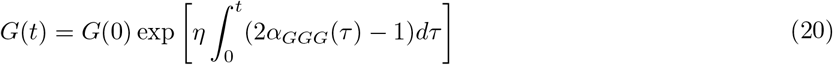

Setting *G*(0) = *P*(0) = 1 (all progenitors are of type *G* at time 0), and plugging into Eq.18, we get:

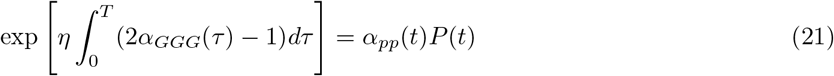

where the variables used in PN model are on the right-hand side (rhs).

At this stage, *α_pp_*(*t*) and *P*(*t*) are in structural correspondence with *α_GGG_* (*t*).

We want to calibrate *α_GGG_* (*t*) from data, so we will use the estimated values of the two former, that we will denote as: *α̂_PP_*(*t*) and *P̂*(*t*).

We seek the *α_GGG_* (*t*) that minimizes the error of prediction upon *α̂_PP_*(*t*) and *P̂*(*t*).

In order to narrow the space of search, we force *α_GGG_* (*t*) to follow the same tanh shape than *α_pp_*(*t*), denoting it:

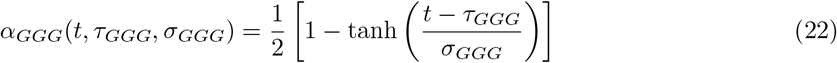

Hence, we then seek the parameters 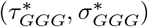 which minimize the square error given by:

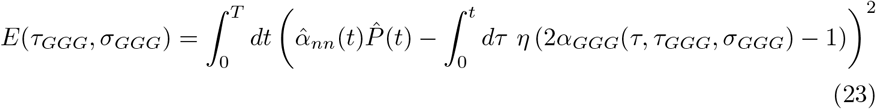

where

- *α̂_nn_*(*t*) is given by Eq. 5 using least-square estimates of *τ_pp_* and *σ_pp_*
- *P̂*(*t*) is given by Eq. 8, using also least-square estimates of *τ_nn_* and *σ_nn_*
- *α_GGG_*(*t*,*τ_GGG_*,*σ_GGG_*) is given by Eq. 22

We note that *τ_GGG_* and *σ_GGG_* are then calibrated using only MoD measures.

We used Nelder-Nead optimization over time-discretized series (with Δ*t* = 0.01 hour, *T* = 96h).

For the three conditions, we found respectively

**CTL :** 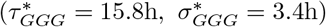

**CDC25B GoF :** 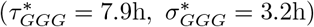

**CDC25B^ΔCDK^ GoF :** 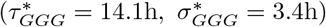

For the record, the corresponding parameters for *τ_pp_* were 23h, 15h and 21h respectively, with *σ_pp_* = 8.0*h*.

We observe that 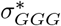 is not affected by the condition, as was *σ_pp_*.

#### 4.2.2 Estimating *G*(*t*)

*G*(*t*) is then solved using Eq. 20.

#### 4.2.3 Estimating *α_GAA_*(*t*)

We simply used:

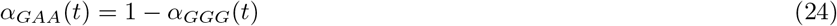

#### 4.2.4 Estimating *α_ANN_*

In the same spirit as for a*GGG*, we should have:

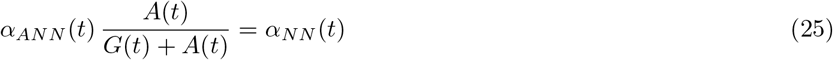

Here again, forcing *α_ANN_*(*t*) to the same tanh shape as *α_NN_*(*t*), we write:

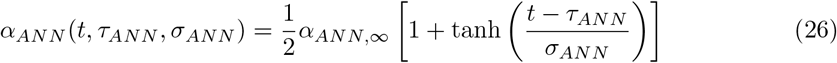

We set *α_ANN,∞_* = 1 as a default value.

Since *G*(*t*) and *α_GAA_* are known from above, we can solve *A*(*t*,*τ_ANN_,σ_ANN_*) for any *α_ANN_*(*t*,*τ_ANN_,σ_ANN_*), using numerical integration to solve :

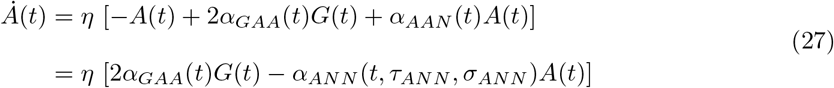

where we used the constraint *α_AAN_* (*t*) + *α_ANN_* (*t*) = 1.

Hence, we seek the best estimates 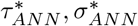 that minimized the square error:

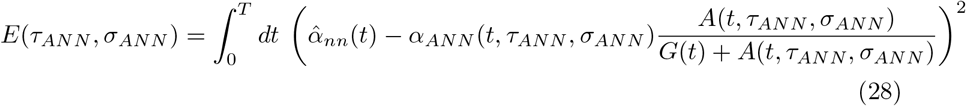

using Nelder-Nead optimization over time-discretized series (with Δ*t* = 0.01 hour, *T* = 96h).

For the three conditions, we found respectively

**CTL :** 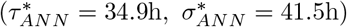

**CDC25B GoF :** 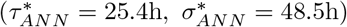

**CDC25B_ΔCDK_ GoF :** 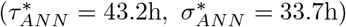

For the record, the corresponding parameters for *α_NN_* are *τ_NN_* = 35.3, 27.3, 39.3 respectively, with *σ_NN_* = 14.5.

#### 4.2.5 Estimating *α_AAN_* (*t*)

We simply used:

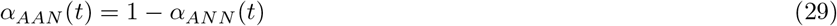

#### 4.2.6 Estimating *A*(*t*)

*A*(*t*) is computed by numerical integration of Eq. 27.

#### 4.2.7 Estimating *N*(*t*)

*N*(*t*) is finally computed by numerical integration of :

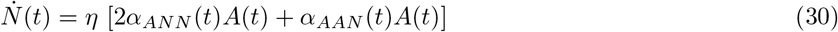

### 4.3 GAG inversion

Estimation of parameters for the GAG model followed the same lines as above, only changing the equations to numerically integrate *G*(*t*), *A*(*t*) and *N*(*t*), namely *Ġ*(*t*), *Ȧ*(*t*), *η*(*t*) equations being changed that have to be changed.

For the three conditions, we found respectively

**CTL :** 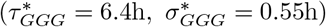

**CDC25B GoF :** 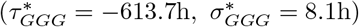

**CDC25B^ΔCDK^ GoF :** 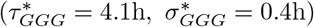

and

**CTL :** 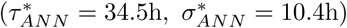

**CDC25B GoF :** 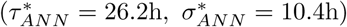

**CDC25B^ΔCDK^ GoF :** 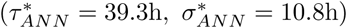

We note that if 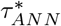 values are pretty similar to those found for GAA model (although not the 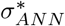 !), the fitting for 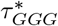 is very bad and yields aberrant values (especially for CDC25B GoF). The good reasons for this are discussed in the text.

## 4.4 GAN inversion

### 4.4.1 Estimating *α_GGG_* (*t*)

Under the *GAN* model, *α_PP_*(*t*)*P*(*t*) only depends upon *α_GGG_* (*t*) so that we can have a more direct expression for it.

At any time *t*:

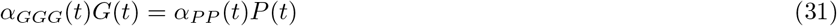

Setting *G*(0) = 1, *G*(*t*) is now a function of *α*_*GGG*_ only:

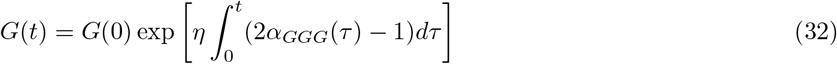

So, in principle, we can calibrate *α_GGG_* directly from Eq.31. We have:

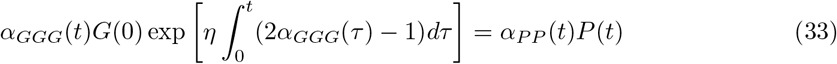

The lhs (left-hand-side) term can be rewritten:

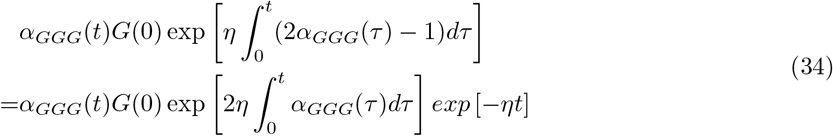

Plugging into Eq.32, and grouping *α_GGG_* terms on the left, we have:

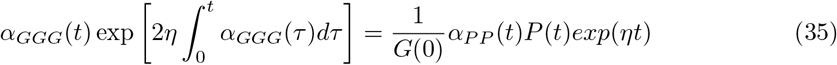

The lhs can be read as a time-derivative:

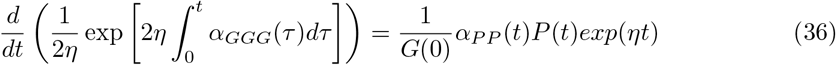

Integrating both sides over [0..*t*] :

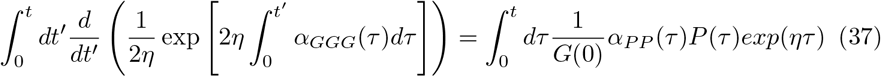

Solving the lhs integral:

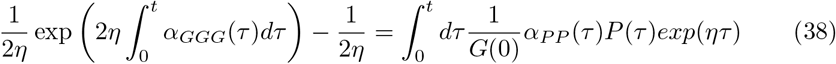

Rearranging terms and taking the ln of both sides :

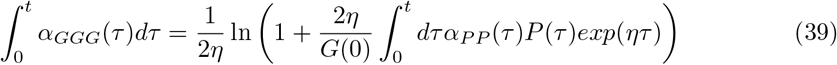

Taking the time derivatives of both sides:

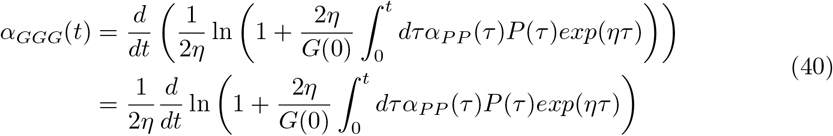

Solving the derivative in the rhs:

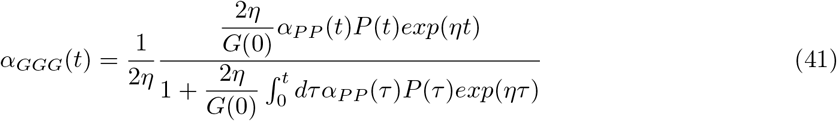

which simplifies to:

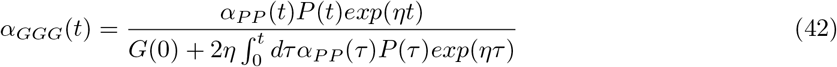

so we take:

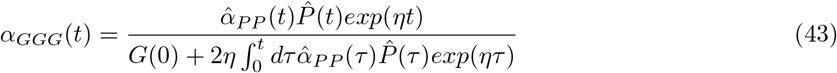

We note that calibrating *α_GGG_*(*t*) by this method does not require to force it to have a tanh shape. In order to check for consistency with the method used to calibrate *α_GGG_*(*t*) under the two other models, we performed the same procedure of minimizing the (corresponding) error function to estimate the 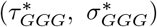.

Both solutions match perfectly, so we retrieve the corresponding parameters 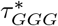 and 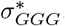.

For the three conditions, we found respectively

**CTL :** 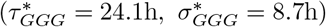

**CDC25B GoF** 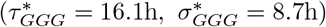

**CDC25B^ΔCDK^ GoF :** 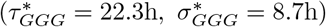

The others parameters were estimated as described for the other models, *Ġ*(*t*), *Ȧ*(*t*), *ṅ*(*t*) equations being changed that have to be changed.

We found:

**CTL :** 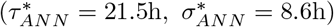

**CDC25B GoF** 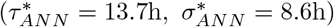

**CDC25B^ΔCDK^ GoF :** 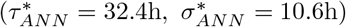

## 5

**List of abbreviations**
BMP: Bone Morphogenetic Protein
CDC25B: Cell division cycle 25 B phosphatase
CDK: Cyclin-dependent kinases
CNS: Central Nervous System
CTL: Control
NSC: Neural Stem Cells
MoD: Mode of Division
Shh: Sonic Hedgehog

## 6 Declarations

### 6.1 Authors’ contributions

All authors contributed to the conceptualisation. MA, SB, JMT and JG contributed to the formal developments. MA, EA and JG wrote the paper. All authors read and approved the final manuscript.

### 6.4 Availability of data and materials

The datasets supporting the conclusions of this article are available upon request to the corresponding author.

